# Universal temporal rate of DNA replication origin firing: A balance between origin activation and passivation

**DOI:** 10.1101/243634

**Authors:** Jean-Michel Arbona, Arach Goldar, Olivier Hyrien, Alain Arneodo, Benjamin Audit

## Abstract

The time-dependent rate *I(t)* of origin firing per length of unreplicated DNA presents a universal bell shape in eukaryotes that has been interpreted as the result of a complex time-evolving interaction between origins and limiting firing factors. Here we show that a normal diffusion of replication fork components towards localized potential replication origins *(p-oris)* can more simply account for the *I(t)* universal bell shape, as a consequence of a competition between the origin firing time and the time needed to replicate DNA separating two neighboring *p-oris*. We predict the *I(t)* maximal value to be the product of the replication fork speed with the squared *p-ori* density. We show that this relation is robustly observed in simulations and in experimental data for several eukaryotes. Our work underlines that fork-component recycling and potential origins localization are sufficient spatial ingredients to explain the universality of DNA replication kinetics.

## Introduction

Eukaryotic DNA replication is a stochastic process (Hyrien et al., 2013; Hawkins et al., 2013; Hyrien, 2016). Prior to entering the S(ynthesis)-phase of the cell cycle, a number of DNA loci called potential origins *(p-oris)* are *licensed* for DNA replication initiation (Machida et al., 2005; Hyrien et al., 2013; Hawkins et al., 2013). During S-phase, in response to the presence of origin *firing* factors, pairs of replication *forks* performing bi-directional DNA synthesis will start from a subset of the *p-oris*, the active replication origins for that cell cycle (Machida et al., 2005; Hyrien et al., 2013; Hawkins et al., 2013). Note that the inactivation of *p-oris* by the passing of a replication fork called origin passivation, forbids origin firing in already replicated regions (de Moura et al., 2010; Hyrien and Goldar, 2010; Yang et al., 2010).

The time-dependent rate of origin firing per length of unreplicated DNA, *I(t)*, is a fundamental parameter of DNA replication kinetics. *I(t)* curves present a universal bell shape in eukaryotes (Goldar et al., 2009), increasing toward a maximum after mid-S-phase and decreasing to zero at the end of S-phase. An increasing *I(t)* results in a tight dispersion of replication ending times, which provides a solution to the random completion problem (Hyrien et al., 2003; Bechhoefer and Marshall, 2007; Yang and Bechhoefer, 2008).

Models of replication in *Xenopus* embryo (Goldar et al., 2008; Gauthier and Bechhoefer, 2009) proposed that the initial *I(t)* increase reflects the progressive import during S-phase of a limiting origin firing factor and its recycling after release upon forks merge. The *I(t)* increase was also reproduced in a simulation of human genome replication timing that used a constant number of firing factors having an increasing reactivity through S-phase (Gindin et al., 2014). In these 3 models, an additional mechanism was required to explain the final *I(t)* decrease by either a subdiffusive motion of the firing factor (Gauthier and Bechhoefer, 2009), a dependency of firing factors’ affinity for *p-oris* on replication fork density (Goldar et al., 2008), or an inhomogeneous firing probability profile (Gindin et al., 2014). Here we show that when taking into account that *p-oris* are distributed at a finite number of localized sites then it is possible to reproduce the universal bell shape of the *I(t)* curves without any additional hypotheses than recycling of fork factor components. *I(t)* increases following an increase of fork mergers, each merger releasing a firing factor that was trapped on DNA. Then *I(t)* decreases due to a competition between the time *t_c_* to fire an origin and the time *t_r_* to replicate DNA separating two neighboring *p-ori*. We will show that when *t_c_* becomes smaller than *t_r_, p-ori* density over unreplicated DNA decreases, and so does *I(t)*. Modeling random localization of active origins in *Xenopus* embryo by assuming that every site is a (weak) *p-ori*, previous work implicitly assumed *t_r_* to be close to zero (Goldar et al., 2008; Gauthier and Bechhoefer, 2009) forbidding the observation of a decreasing *I(t)*. Licensing of a limited number of sites as *p-ori* thus appears to be a critical property contributing to the observed canceling of *I(t)* at the end of S-phase in all studied eukaryotes.

## Results

### Emergence of a bell-shape *I(t)*

In our modeling of replication kinetics, a bimolecular reaction between a firing factor and a *p-ori* results in an origin firing event; then the diffusing element is trapped and travels with the replication forks until two converging forks merge (termination, Fig. 1 (a)). Under the assumption of a well-mixed system, for every time step *dt*, we consider each interaction between the *N_FD_(t)* free diffusing firing factors and the *N_p-ori_(t) p-oris* as potentially leading to a firing with a probability *k_on_dt*. The resulting simulated firing rate per length of unreplicated DNA is then:

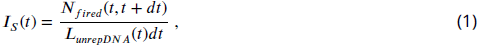

where *N_fired_(t),t + dt)* is the number of *p-oris* fired between times *t* and *t + dt*, and *L_unrepDNA_(t)* is the length of unreplicated DNA a time *t*. Then we propagate the forks along the chromosome with a constant speed *v*, and if two forks meet, a free firing factor is released. Finally we simulate the chromosomes as 1D chains where prior to entering S-phase, the *p-oris* are precisely localized. For *Xenopus* embryo, the *p-ori* positions are randomly sampled, so that each simulated S-phase corresponds to a different positioning of the *p-oris*. We compare results obtained with periodic or uniform *p-ori* distributions. For *S. cerevisiae*, the *p-ori* positions, identical for each simulation, are taken from the OriDB database (Siow et al., 2012). As previously simulated in human (Löb et al., 2016), we model the entry in S-phase using an exponentially relaxed loading of the firing factors with a time scale shorter than the S-phase duration *T_phase_* (3 mins for *Xenopus* embryo, where *T_phase_* ~ 30 mins, and 10 mins for *S. cerevisiae*, where *T_phase_* ~ 60 mins). After the short loading time, the total number of firing factors 
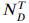
 is constant. As shown in Fig. 1 (b) (see also Fig. 2), the universal bell shape of the *I(t)* curves (Goldar et al., 2009) spontaneously emerges from our model when going from weak to strong interaction, and decreasing the number of firing factors below the number of *p-oris*. The details of the firing factor loading dynamics do not affect the emergence of a bell shaped *I(t)*, even though it can modulate its precise shape, especially early in S-phase.

In a simple bimolecular context, the rate of origin firing is *i(t) = k_on_N_p-ori_(t)N_FD_(t)*. The firing rate by element of unreplicated DNA is then given by

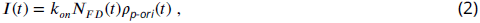

where *ρ_p-ori_(t) = N_p-ori_(t)/L_unrepDNA_(t)*. In the case of a strong interaction and a limited number of firing factors, all the diffusing factors react rapidly after loading and *N_FD_(t)* is small (Fig. 1 (c), dashed curves). Then follows a stationary phase where as long as the number of *p-oris* is high (Fig. 1 (c), solid curves), once a diffusing factor is released by the encounter of two forks, it reacts rapidly, and so *N_FD_(t)* stays small. Then, when the rate of fork mergers increases due to the fact that there are as many active forks but a smaller length of unreplicated DNA, the number of free firing factors increases up to 
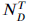
 at the end of S-phase. As a consequence, the contribution of *N_FD_(t)* to *I(t)* in Eq. (2) can only account for a monotonous increase during the S phase. For *I(t)* to reach a maximum *I_max_* before the end of S-phase, we thus need that *ρ_p-ori_(t)* decreases in the late S-phase. This happens if the time to fire a *p-ori* is shorter than the time to replicate a typical distance between two neighboring *p-oris*. The characteristic time to fire a *p-ori* is *t_c_ = 1/k_on_N_FD_(t)*. The mean time for a fork to replicate DNA between two neighboring *p-oris* is *t_r_ = d(t)/v*, where *d(t)* is the mean distance between unreplicated *p-oris* at time *t*. So the density of origins is constant as long as:

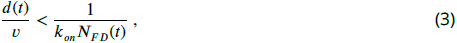

or

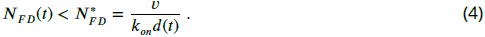

**Figure 1.**
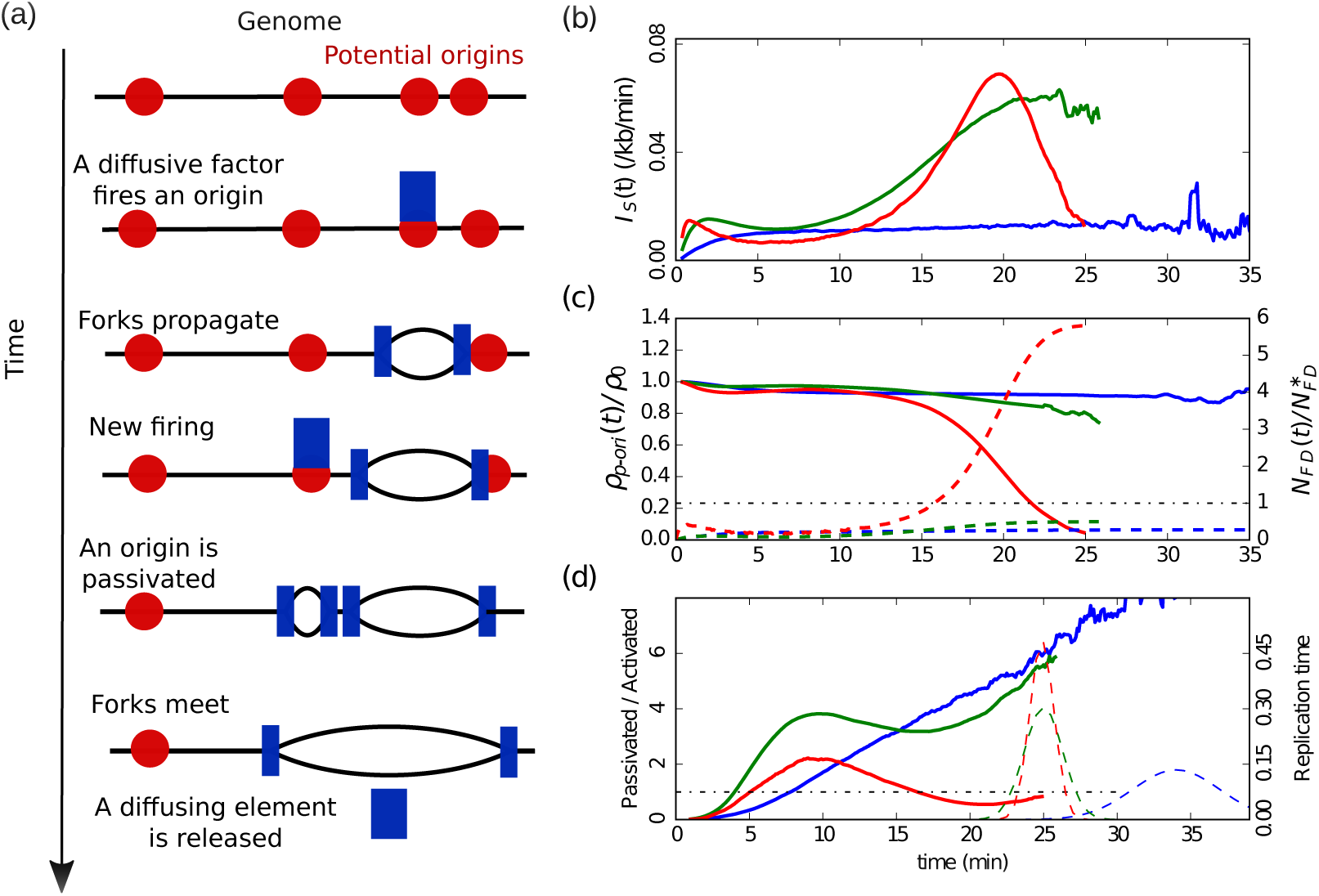
(a) Sketch of the different steps of our modeling of replication initiation and propagation. (b) *IS_(t)_* (Eq. (1)) obtained from numerical simulations of one chromosome of length 3000 kb, with a fork speed *v* = 0.6 kb/min. The firing factors are loaded with a characteristic time of 3 mins. From blue to green to red the interaction is increased and the number of firing factors is decreased: blue (*k_on_* = 5+10^−5^ min^−1^, 
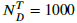
, *ρ*_0_ = 0.3 kb^−1^), green (*k_on_* = 6+10^−4^ min^−1^, 
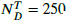
, *ρ*_0_ = 0.5 kb^−1^), red (*k_on_* = 6+10^−3^ min^−1^, 
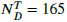
, *ρ*_0_ = 0.28 kb^−1^)). (c) Corresponding normalized densities of *p-oris* (solid lines), and corresponding normalized numbers of free diffusing firing factors (dashed line): blue 
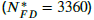
, green 
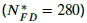
, red 
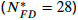
; the light blue horizontal dashed line corresponds to the critical threshold value 
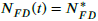
. (d) Corresponding number of passivated origins over the number of activated origins (solid lines). Corresponding histograms of replication time (dashed lines).

Thus, at the beginning of the S-phase, *N_FD_(t)* is small, *ρ_p-ori_>(t)* is constant (Fig. 1 (c), solid curves) and so *I_S_(t)* stays small. When *_NFD_(t)* starts increasing, as long as Eq. (4) stays valid, *I_S_(t)* keeps increasing. When *_NFD_(t)* becomes too large and exceeds 
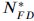
 then Eq. (4) is violated and the number of *p-oris* decreases at a higher rate than the length of unreplicated DNA, and *ρ_p-ori_(t)* decreases and goes to zero (Fig. 1 (c), red solid curve). As *_NFD_(t)* tends to 
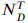
 *I_S_(t)* goes to zero, and its global behavior is a bell shape (Fig. 1 (b), red).

Let us note that if we decrease the interaction strength (*k_on_*), then the critical 
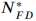
 will increase beyond 
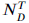
 (Fig. 1 (c), dashed blue and green curves). *I_S_(t)* then monotonously increase to reach a plateau (Fig. 1 (b), green), or if we decrease further *k_on_, I_S_(t)* present a very slow increasing behavior during the S-phase (Fig. 1 (b), blue). Now if we come back to strong interactions and increase the number of firing factors, almost all the *p-oris* are fired immediately and *I_S_(t)* drops to zero after firing the last *p-ori*.

Another way to look at the density of *p-oris* is to compute the ratio of the number of passivated origins by the number of activated origins (Fig. 1 (d)). After the initial loading of firing factors, this ratio is higher than one. for weak and moderate interactions (Fig. 1 (d), blue and green solid curves, respectively) this ratio stays bigger than one during all the S-phase, where *I_S_(t)* was shown to be monotonously increasing (Fig. 1 (b)). For a strong interaction (Fig. 1 (b), red solid curve), this ratio reaches a maximum and then decreases below one, at a time corresponding to the maximum observed in *I_S_(t)* (Fig. 1 (d), red solid curve). Hence, the maximum of *I(t)* corresponds to a switch of the balance between origin passivation and activation, the latter becoming predominant in late S-phase. We have seen that up to this maximum *ρ_p-ori_(t) ≈ cte ≈ ρ_0_*, so *I_S_(t) ≈ k_on_ ρ_0_ N_F_(t)*. When *N_FD_(t)* reaches 
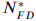
 then *I_S_(t)* reaches its maximum value:

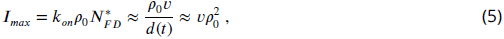

where we have used the approximation *d(t) ≈ d(0) = 1/ρ_0_* (which is exact for periodically distributed *p-oris*). *I_max_* can thus be predicted from two measurable parameters, providing a direct test of the model.

### Comparison with different eukaryotes

*Xenopus* embryo. Given the huge size of *Xenopus* embryo chromosomes, to make the simulations more easily tractable, we rescaled the size *L* of the chromosomes, *k_on_* and 
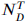
 to keep the duration of S-phase 
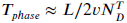
 and *I(t)* (Eq. (2)) unchanged 
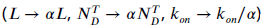
. In Fig. 2 (a) are reported the results of our simulations for a chromosome length *L* = 3000 kb. We see that a good agreement is obtained with experimental data (Goldar et al., 2009) when using either a uniform distribution of *p-oris* with a density *ρ*_0_ = 0.70 kb^−1^ and a number of firing factors 
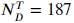
, or a periodic distribution with *ρ*_0_ = 0.28 kb^−1^ and 
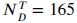
. A higher density of *p-oris* was needed for uniformly distributed *p-oris* where *d(t)* (slightly) increases with time, than for periodically distributed *p-oris* where *d(t)* fluctuates around a constant value 1/*ρ*_0_. The uniform distribution, which is the most natural to simulate *Xenopus* embryo replication, gives a density of activated origins of 0.17 kb^−1^ in good agreement with DNA combing data analysis (Herrick et al., 2002) but twice lower than estimated from real time replication imaging of surface-immobilized DNA in a soluble *Xenopus* egg extract system (Loveland et al., 2012).

*S. cerevisiae*. To test the robustness of our minimal model with respect to the distribution of *p-oris*, we simulated the replication in *S. cerevisiae*, whose *p-oris* are known to be well positioned as reported in OriDB (Siow et al., 2012). 829 *p-oris* were experimentally identi1ed and classi1ed into three categories: confirmed origins (410), likely origins (216), and dubious origins (203). When comparing the results obtained with our model to the experimental *I(t)* data (Goldar et al., 2009) (Fig. 2 (b)), we see that to obtain a good agreement we need to consider not only the confirmed origins but also the likely and the dubious origins. However in regard to the uncertainty in the value of the replication fork velocity and the possible experimental contribution of the *p-oris* in the rDNA part of chromosome 12 (not taken into account in our modeling), this conclusion needs to be confirmed in future experiments. It is to be noted that even if 829 *p-oris* are needed, on average only 352 origins have fired by the end of S-phase. For *S. cerevisiae* with well positioned *p-oris*, we have checked the robustness of our results with respect to a stochastic number of firing factors 
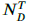
 from cell to cell (Poisson distribution, Iyer-Biswas et al. (2009)). We confirmed the *I(t)* bell shape with a robust duration of the S-phase of 58.6±4.3 min as compared to 58.5±3.3 min obtained previously with a constant number of firing factors. Interestingly, in an experiment where *T_phase_* was lengthened from 1 h to 16 h by adding hydroxyurea (HU) in yeast growth media, the pattern of activation of replication origins was shown to be conserved (Alvino et al., 2007). HU slows down the DNA synthesis to a rate of ~ 50 bp min^−1^ corresponding to a 30 fold decrease of the fork speed (Sogo et al., 2002). In our model with a constant number of firing factors, 
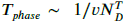
 a two fold increase of the number 
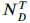
 of firing factors is sufficient to account for the 16 fold increase of *T_phase_*, which is thus mainly explained by the HU induced slowdown of the replication forks. In a model where the increase of *I(t)* results from the import of replication factors, the import rate would need to be reduced by the presence of HU in proportion with the lengthening of S-phase in order to maintain the pattern of origin activations. Extracting *I(t)* from experimental replication data for cells grown in absence (HU^−^) or presence (HU^+^) (Alvino), we estimated 
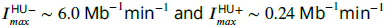
 for HU^−^ and HU^+^ cells, respectively. The ratio 
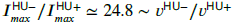
 is quite consistent with the prediction of the scaling law (Eq. (5)) for a constant density of *p-oris*.

**Figure 2.**
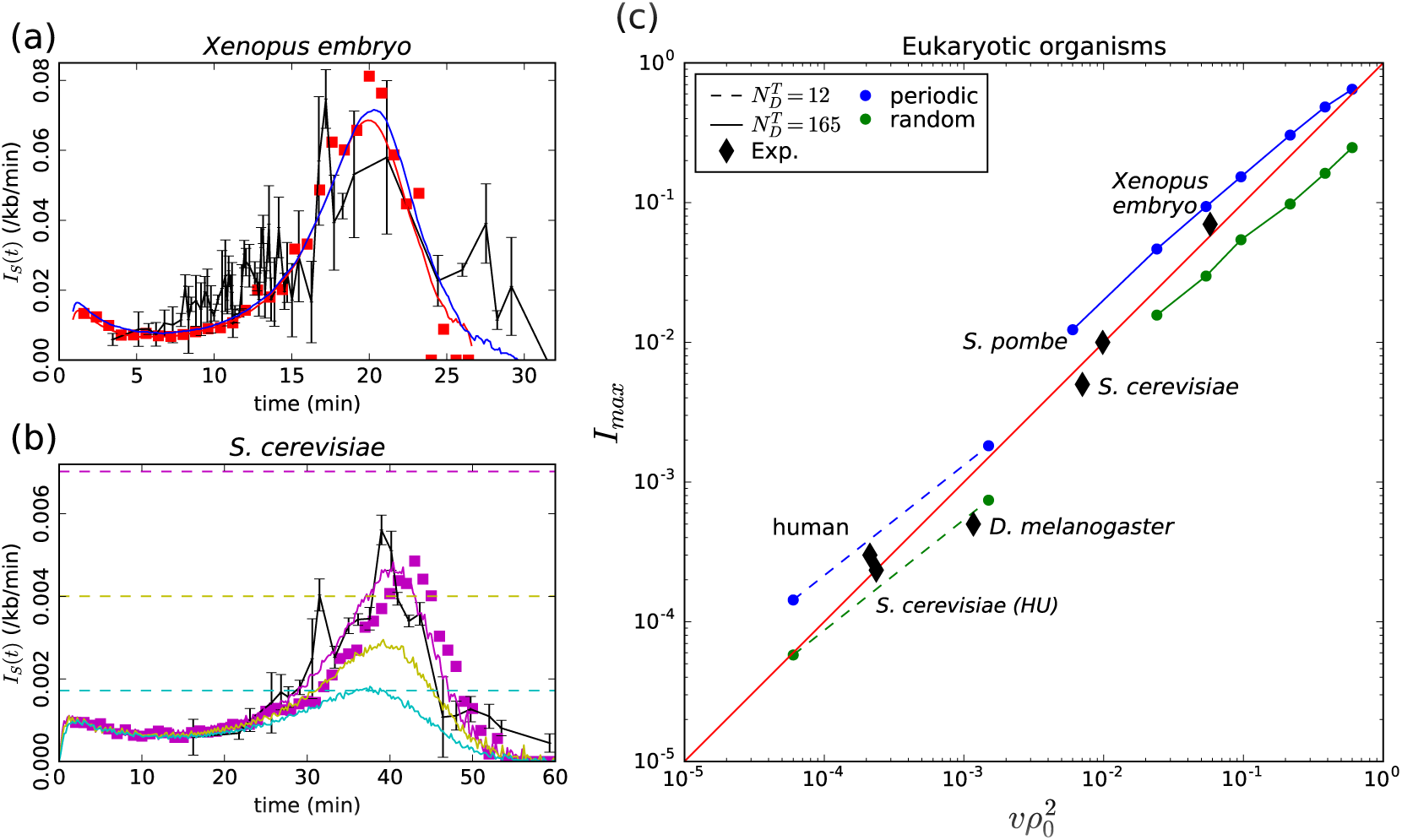
(a) *Xenopus* embryo: Simulated *I_S_(t)* (Eq. (1)) for a chromosome of length *L* = 3000 kb and a uniform distribution of *p-oris* (blue: *v* = 0.6 kb/min, *k_on_* = 3.+10^−3^ min^−1^, 
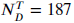
 *ρ*_0_ = 0.70 kb^−1^) or a periodic distribution of *p-oris* (red: *v* = 0.6 kb/min, *k_on_* = 6×10^−3^ min^−1^ 
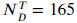
 *ρ*_0_ = 0.28 kb^−1^); (red squares) 3D simulations with the same parameter values as for periodic *p-ori* distribution; (black) experimental *I(t)*: raw data obtained from Goldar et al. (2009) were binned in groups of 4 data points; the mean value and standard error of the mean of each bin were represented. (b) *S. cerevisiae*: Simulated *I_S_(t)* for the 16 chromosomes with the following parameter values: *v* = 1.5 kb/min, 
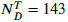
 *k_on_* = 3.6×10^−3^ min^−1^, when considering only confirmed origins (light blue), confirmed and likely origins (yellow) and confirmed, likely and dubious origins (purple); the horizontal dashed lines mark the corresponding predictions for *I_max_* (Eq. (5)); (purple squares) 3D simulations with the same parameter values considering confirmed, likely and dubious origins; (black) experimental *I(t)* from Goldar et al. (2009). (c) Eukaryotic organisms: *I_max_* as a function of 
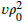
 (squares and bullets) simulations performed for regularly spaced origins (blue) and uniformly distributed origins (green) with two sets of parameter values: *L* = 3000 kb, *v* = 0.6 kb/min, *k_on_* = 1.2×10^−2^ min^−1^ and 
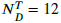
 (dashed line) or 165 (solid line); (black diamonds) experimental data points for *Xenopus* embryo, *S. cerevisiae*, *S. cerevisae* grown in Hydroxyurea (HU), *S. pombe, D. melanogaster*, human (see text and Table 1).

**Table 1.** Experimental data for various eukaryotic organisms with genome length *L (Mb)*, replication fork velocity *v* (kb/min), number of *p-oris (N_p-ori_(t=0)), ρ_0_ = N_p-ori_(t=0)/L (kb^−1^)* and *I_max_ (Mb^−1^min^−1^)*. All *I_max_* data are from Goldar et al. (2009), except for *S. cerevisiae* grown in presence or absence of hydroxyurea (HU) which were computed from the replication profile of Alvino et al. (2007). For *S. cerevisiae* and *S. pombe*, confirmed, likely, and dubious origins were taken into account. For *D. melanogaster*, *N_p-ori_(t=0)* was obtained from the same Kc cell type as the one used to estimate *I_max_*. For *Xenopus* embryo, we used the experimental density of activated origins to estimate *N_p-ori_(t=0)* which is probably lower than the true number of *p-oris*. For human, we averaged the number of origins experimentally identi1ed in K562 (62971) and in MCF7 (94195) cell lines.

| | $L$ | $v$ | $N_{p-ori}$ | $\rho_0$ | $I_{max}$ | Ref. |
| --- | --- | --- | --- | --- | --- | --- |
| *S. cerevisiae* | 12.5 | 1.60 | 829 | 0.066 | 6.0 | Sekedat et al. (2010); Siow et al. (2012) |
| *S. cerevisiae* in presence of HU | 12.5 | 0.05 | 829 | 0.066 | 0.24 | Alvino et al. (2007). Same $N_{p-ori}$ and $\rho_0$ as *S. cerevisiae* in normal growth condition. |
| *S. pombe* | 12.5 | 2.80 | 741 | 0.059 | 10.0 | Siow et al. (2012); Kaykov and Nurse (2015) |
| *D. melanogaster* | 143.6 | 0.63 | 6184 | 0.043 | 0.5 | Ananiev et al. (1977); Cayrou et al. (2011) |
| *Xenopus* embryo | 2233.0 | 0.52 | 744333 | 0.333 | 70.0 | Loveland et al. (2012) |
| human | 6469.0 | 1.46 | 78000 | 0.012 | 0.3 | Conti et al. (2007); Martin et al. (2011) |

*D. melanogaster* and human. We gathered from the literature experimental estimates of *I_max_, ρ_0_* and *v* for different eukaryotic organisms (Table 1). As shown in Fig. 2 (c), when plotting 
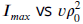
 all the experimental data points remarkably follow the diagonal trend indicating the validity of the scaling law (Eq. (5)) for all considered eukaryotes. We performed two series of simulations for fixed values of parameters *k_o_*, 
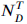
 and *v* and decreasing values of *ρ*_0_ with both periodic distribution (blue) and uniform (green) dist tions of *p-oris* (Fig. 2 (c)). The first set of parameters was chosen to cover high *I_max_* values similar the one observed for *Xenopus* embryo (bullets, solid lines). When decreasing *ρ*_0_, the number of firing factors becomes too large and *I(t)* does no more present a maximum. We thus decreased the value of 
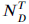
 keeping all other parameters constant (boxes, dashed line) to explore smaller values of Imax in the range of those observed for human and *D. melanogaster*. We can observe that experimental data points’ deviation from Eq. (5) is smaller than the deviation due to specific *p-oris* distributions.

## Discussion

To summarize, we have shown that within the framework of 1D nucleation and growth models of DNA replication kinetics (Herrick et al., 2002; Jun and Bechhoefer, 2005), the sufficient conditions to obtain a universal bell shaped *I(t)* as observed in eukaryotes are a strong bimolecular reaction between localized *p-oris* and limiting origin firing factors that travel with replication forks and are released at termination. Under these conditions, the density of *p-oris* naturally decreases by the end of the S-phase and so does *I_S_(t)*. Previous models in *Xenopus* embryo (Goldar et al., 2008; Gauthier and Bechhoefer, 2009) assumed that all sites contained a *p-ori* implying that the time *t_r_* to replicate DNA between two neighboring *p-oris* was close to zero. This clari1es why they needed some additional mechanisms to explain the final decrease of the firing rate. Moreover our model predicts that the maximum value for *I(t)* is intimately related to the density of *p-oris* and the fork speed (Eq. (5)), and we have shown that without free parameter, this relationship holds for 5 species up to a 300 fold difference of *I_max_* and 
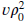
 (Table 1, Fig. 2 (c)).

In contrast with models where replication kinetics is explained by properties specific to each *p-oris* (Bechhoefer and Rhind, 2012), our model assumes that all *p-oris* are governed by the same rule of initiation resulting from physicochemically realistic particulars of their interaction with limiting replication firing factors. To confirm this simple physical basis of our modeling, we used molecular dynamics rules as previously developed for *S. cerevisiae* (Arbona et al., 2017) to simulate S-phase dynamics of chromosomes confined in a spherical nucleus. We added firing factors that are free to diffuse in the covolume left by the chain and that can bind to proximal *p-oris* to initiate replication, move along the chromosomes with the replication forks and be released when two fork merges. As shown in Fig. 2 (a, b) for *Xenopus* embryo and *S. cerevisiae*, results confirmed the physical relevance of our minimal modeling and the validity of its predictions when the 3D diffusion of the firing factors is explicitly taken into account. This opens new perspectives for understanding correlations between firing events along chromosomes that could result in part from the spatial transport of firing factors. For example in *S. cerevisiae* (Knott et al., 2012) and in *S. pombe* (Kaykov and Nurse, 2015), a higher firing rate has been reported near origins that have just fired (but see Yang et al. (2010)). In mammals, megabase chromosomal regions of synchronous firing were first observed long ago (Huberman and Riggs, 1968; Hyrien, 2016). Recently, profiling of replication fork directionality obtained by Okazaki fragment sequencing have suggested that early firing origins located at the border of Topologically Associating Domains (TADs) trigger a cascade of secondary initiation events propagating through the TAD (Petryk et al., 2016). Early and late replicating domains were associated with nuclear compartments of open and closed chromatin (Ryba et al., 2010; Boulos et al., 2015; Goldar et al., 2016; Hyrien, 2016). In human, replication timing U-domains (0.1-3 Mb) were shown to correlate with chromosome structural domains (Baker et al., 2012; Moindrot et al., 2012; Pope et al., 2014) and chromatin loops (Boulos et al., 2013, 2014).

Understanding to which extent spatio-temporal correlations of the replication program can be explained by the diffusion of firing factors in the tertiary chromatin structure specific to each eukaryotic organism is a challenging issue for future work.

We thank F. Argoul for helpful discussions. This work was supported by Institut National du Cancer (PLBIO16-302), Fondation pour la Recherche Médicale (DEI20151234404) and Agence National de la Recherche (ANR-15-CE12-0011-01). BA acknowledges support from Science and Technology Commission of Shanghai Municipality (15520711500) and Joint Research Institute for Science and Society (JoRISS). We gratefully acknowledge support from the PSMN (Pôle Scientifique de Modélisation Numérique) of the ENS de Lyon for the computing resources. We thank BioSyL Federation and Ecofect LabEx (ANR-11-LABX-0048) for inspiring scienti1c events.

